# Biomechanical characterisation of thoracic ascending aorta with preserved pre-stresses

**DOI:** 10.1101/2022.12.02.518810

**Authors:** Shaiv Parikh, Kevin M. Moerman, Mitch J.F.G. Ramaekers, Simon Schalla, Elham Bidar, Tammo Delhaas, Koen Reesink, Wouter Huberts

## Abstract

Mechanical properties of an aneurysmatic thoracic aorta are potential markers of future growth and remodeling and can help to estimate risk of rupture. Aortic geometries obtained from routine medical imaging do not display wall stress distribution and mechanical properties. Mechanical properties for a given vessel may be determined from medical images at different physiological pressures using inverse finite element analysis. However, without considering pre-stresses, the estimation of mechanical properties will lack accuracy. In the present paper, we propose and evaluate a mechanical parameter identification technique, which recovers pre-stresses by determining the zero-pressure configuration of the aortic geometry. We first validated the method on a cylindrical geometry and subsequently applied it to a realistic aortic geometry. Verification of the assessed parameters was performed using synthetically generated reference data for both geometries. The method was able to estimate the true mechanical properties with an accuracy ranging from 98% to 99%.

**Author summary:** Current clinical guidelines recommend to perform surgical intervention on thoracic ascending aortas with diameters greater than 55 mm. However, diameter sizes solely are not indicative of the degenerative changes in the vessel wall. Patient-specific tissue mechanical properties give better insight into the degenerative changes caused due to the variations in mass densities of the vessel wall constituents. Inverse finite element analysis can be performed on aortic geometries derived from medical images to determine tissue mechanical properties. Performing inverse finite element analysis on geometries obtained directly from medical images leads to incorrect estimation of vessel wall stresses and therefore, mechanical properties. Mechanical properties estimation is incorrect because inverse analysis does not take into account the pre-stressed nature of the geometry. In this study, we propose a novel approach to non-invasively estimate *in vivo* mechanical properties of the thoracic ascending aorta by performing inverse finite element analysis, which considers the vessel wall pre-stresses.

## Introduction

Aortic aneurysms are localized dilations, characterised by degeneration of the vessel wall [1, 2]. If left untreated, aneurysms may rupture or dissect, thereby leading to high mortality rates [3]. The main load bearing constituents of the aortic wall are elastin, collagen fibers and vascular smooth muscle cells (VSMCs) [1].Variations in concentrations, reorientation, or change in mechanical properties of these constituents reflect changes in mechanical properties of the tissue [1]. Eventual rupture of an aneurysm is observed when the aortic wall material fails under certain physiological forces exerted on it [1, 2]. It has been shown that stiffening of aortic wall, e.g. by loss of elastin, is associated with enlargement of the aneurysm [1, 2]. Therefore, estimation of aortic mechanical properties might prove crucial for clinicians to understand the stage of aneurysm development or to determine the risk of aneurysm rupture or dissection.

Use of inverse finite element analysis (IFEA) is a common approach in biomechanics for non-invasive assessment of mechanical properties [4]. IFEA typically involves creating a finite element (FE) geometry of the object of interest, assuming a constitutive law with initial mechanical property parameters, applying realistic loads to the FE geometry and computing the outcome in terms of motion and deformation. The computed outcomes are then compared to *in vivo* experimental data, such as measured deformations. The mechanical parameters are iteratively modified until best-fit estimates are obtained [4].

Accurate creation of FE models requires accurate aortic geometries. These can be obtained through segmentation of medical image data obtained from computed tomography (CT), magnetic resonance imaging (MRI) or ultrasound [5, 6]. However, these image-based geometries cannot immediately be used for a finite element analysis as they are acquired under *in vivo* loading conditions with a pre-existing strain and hence, a stress field in the vessel wall [5, 6]. The stress field present under *in vivo* conditions is due to the residual stresses as well as pre-stresses. Residual stresses are defined as stresses arising in the aortic wall due to different stretches that the vessel constituents (elastin, collagen) experience individually under zero-pressure configuration. Pre-stresses are defined as the stresses arising in the aortic wall by pressurising the zero-pressure unloaded configuration [5, 6]. Commonly, the geometry obtained from the medical images is tacitly accepted as zero-stress configuration but this will result in incorrect values of stresses in the vessel wall during IFEA, and consequently incorrect parameter estimation [5, 6].

*In vivo* assessment of residual stresses is not possible without a priori knowledge of tissue mechanical properties [7]. Nevertheless, it can be assumed that the effect of residual stresses is lower than the effect of pre-stresses, and therefore, the effect of residual stresses can be neglected [6, 8, 9]. Determination of pre-stresses remains imperative and is possible by estimating the unloaded configuration using methods such as – (i) determination of zero-pressure configuration through a minimisation problem, (ii) the backward displacement, and (ii) the backward incremental method [5, 6, 8]. The former two methods deal with the fact that the determination of the unloaded configuration is dependent on the a priori knowledge of mechanical parameters of the geometry [5, 8]. Though the backward incremental method has demonstrated its ability to estimate mechanical parameters as well as pre-stresses [6, 9], it is not straightforward to use this method since, the FE code of the structural solver needs to be adapted to update the stress tensor field at each pressure increment without updating the deformation field to build up the pre-stresses [5].

The aim of the present study was to formulate a method for IFEA-based non-invasive mechanical property identification, including proper consideration of pre-stresses, and not requiring alterations to the FE solver. Our method incorporates elements from previous methods to determine zero-pressure geometries non-invasively [5, 8], while simultaneously estimating mechanical properties of the ascending aorta. For validation, we applied our method to synthetically generated reference data of a simple cylindrical geometry first and subsequently to that of a complex, real aortic geometry.

## Methods

The focus of this paper is to validate a mechanical parameter estimation algorithm for a general aortic geometry. The validation of the algorithm is performed by implementing the workflow on synthetically derived reference data as a substitute for patient-derived experimental data. Therefore, to understand the validation process, the algorithm must be read by replacing “experimental data” with “synthetic reference data”.

The key elements required for the mechanical parameter estimation entail geometry creation (in this paper: cylindrical geometry and an image-based aortic geometry), a FE model and an inverse modelling algorithm that considers existing pre-stresses. Each of these components are described in this section. At the end of the section, the creation of synthetic reference data and the simulations that were performed to assess the accuracy of our IFEA are presented.

### Vessel Geometries

In this section, the geometry creation of our two computational domains, i.e., cylinder and aorta, will be explained. Additionally, the boundaries where pressures and displacements are applied, are also explained. All operations of geometry creation were performed using MATLAB codes developed based on the open source GIBBON toolbox [10].

### Cylindrical geometry

A cylindrical geometry was created with a radius of 16 mm and a length of 192 mm. Subsequently, the geometry was meshed using quadrilateral elements. The mesh was thickened by 1 mm on the inner side to create an 8-noded hexahedral element mesh. This resulted in inner radius of 15 mm for the entire geometry. The meshed geometry, after checking for mesh independency, comprised of 2176 elements with 4420 nodes denoted as 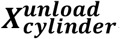. The nodes on the openings of the cylinder (*Γ*_*d*_; cf. Fig 1) were provided with fixed boundary conditions during all simulations, while the inner surface of the cylinder (*Γ*_*n*_; cf. Fig 1) was prescribed with pressure load.

**Fig 1.**
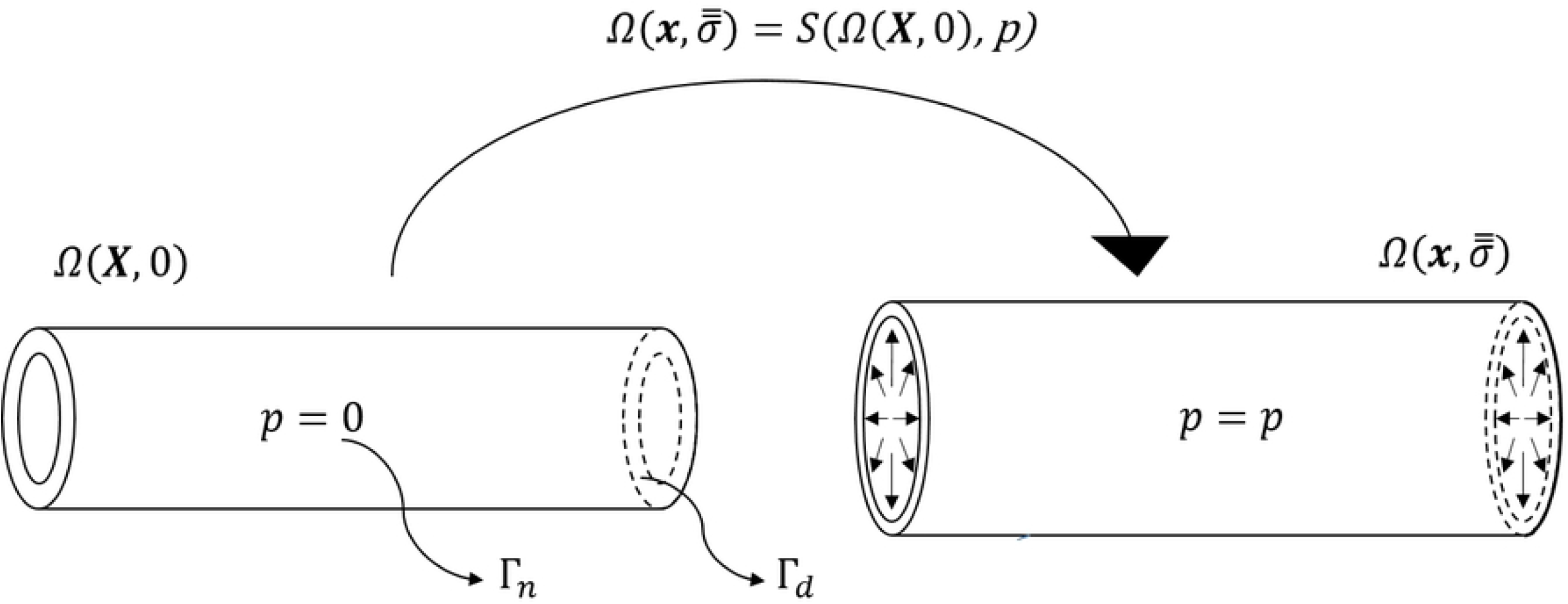
Schematic representation of computational domain, *Ω*, of the cylindrical geometry. The finite element system of equations are solved by a structural solver (*S*) by applying prescribed displacements at the opening boundaries (*Γ*_*d*_) and pressure (*p*) at the inner surface (*Γ*_*n*_). The general notation for nodes in the undeformed reference configuration and the deformed configuration are represented as ***X*** and ***x*** respectively, with stresses in the deformed configuration as 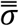

### Image-based aorta geometry

The aortic geometry at diastolic pressure was reconstructed from DICOM data of the 3D DIXON sequence generated from 3-T MR systems (Philips Ingenia; Philips Healthcare, Best, the Netherlands). From the DICOM data, *x, y* and *z* coordinates of four points around the luminal edges were retrieved for six anatomical locations (Fig 2A).

**Fig 2.**
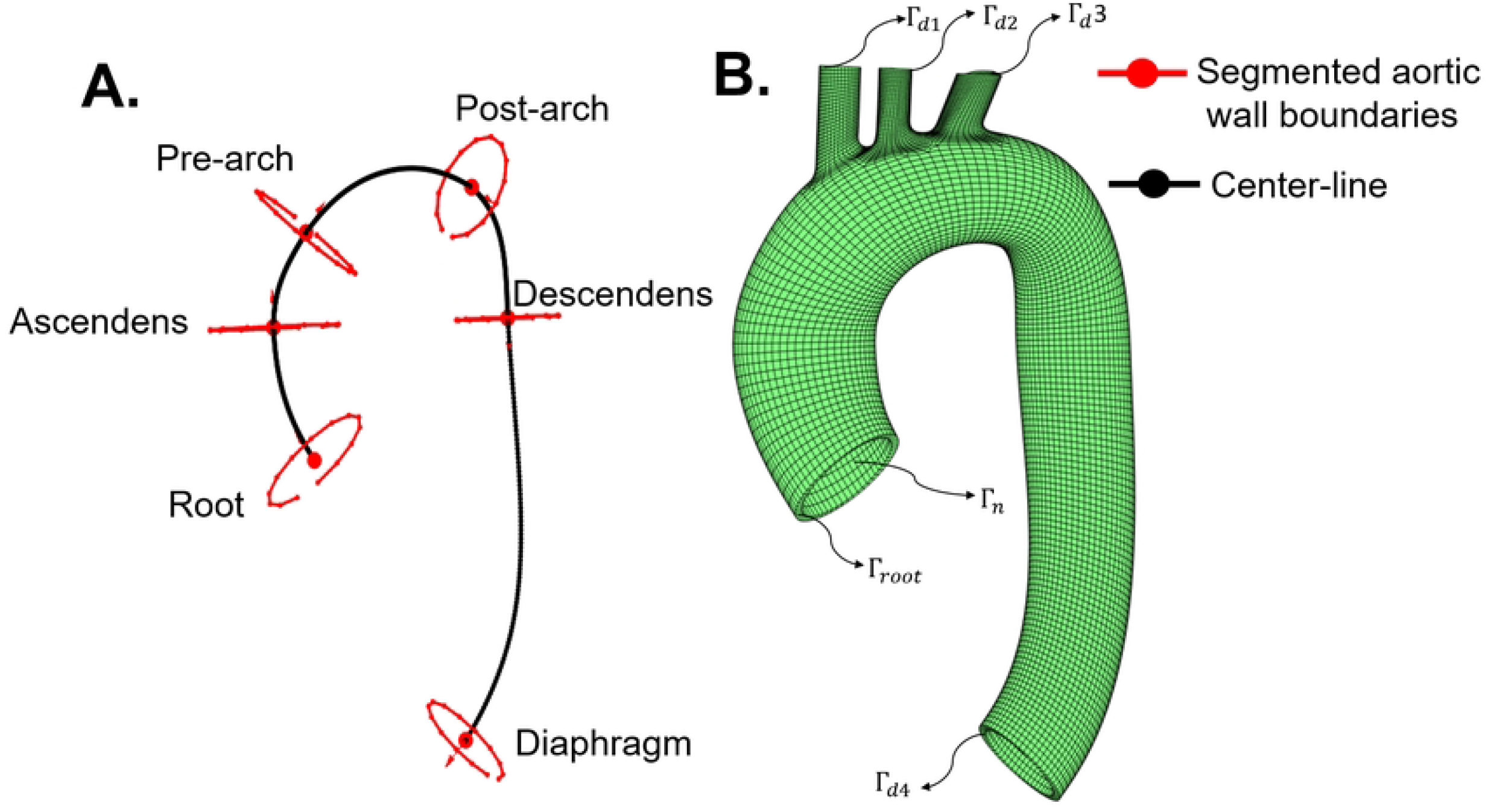
(A) Inner-wall aortic boundaries obtained from 3D DIXON MRI scan and centre-line connecting the lumen at six plane locations. The six plane locations were at root, ascending aorta, before brachiocephalic artery (pre-arch), after left sub-clavian artery (post-arch), descending aorta, and at diaphragm levels. (B) Solid mesh (*Ω* (***X***^***MRI***^,0)) of the aortic geometry including the branches. Boundary conditions were applied on the nodes located on *Γ*_*d*1_ to *Γ*_*d*4_ (fixed), *Γ*_*root*_ (fixed/displacement), and *Γ*_*n*_ (pressure).

Subsequently, smooth ellipses were fitted through the retrieved four points from the luminal edges. The outer boundaries were then created by adding a layer of points at an offset of 2mm from the inner boundaries (at each plane location). A curve connecting the centroids of the outer-wall boundaries through a cubic polynomial fit represented the centre-line of the main trunk of the aorta. Similar to the cylindrical model, a quadrilateral surface-mesh was swept along the centre-line of the main trunk. Next, major branch vessels were added. The mesh was thickened in the inwards direction (2mm thickness) to create 8-noded hexahedral elements (Fig. 2B). The geometry resulted in 11998 elements with 24236 nodes denoted as ***X***^***MRI***^. All boundary conditions were applied on the unloaded configuration (detailed explanation in next section). The nodes on openings (*Γ*_*d*1_ to *Γ*_*d*4_; Fig 2B) were prescribed with fixed (Dirichlet) boundary conditions while pressurising the inner surface *Γ*_*n*_ (Fig 2B) of the unloaded configuration to diastolic pressure. However, the nodes at the root level *Γ*_*root*_ (Fig 3B) were prescribed with displacement while pressurising the unloaded configuration to systolic pressure to take into account the longitudinal motion of the root during a cardiac cycle. The root movement was determined as follows: the aortic valve movement was derived from single slice cine images of the left ventricular outflow tract (LVOT), which was acquired in two directions (perpendicular to each other). Valve displacement was measured automatically by using commercially available software (CAAS MR Solutions v 5.2.1, PieMedical Imaging, Maastricht, the Netherlands).

**Fig 3.**
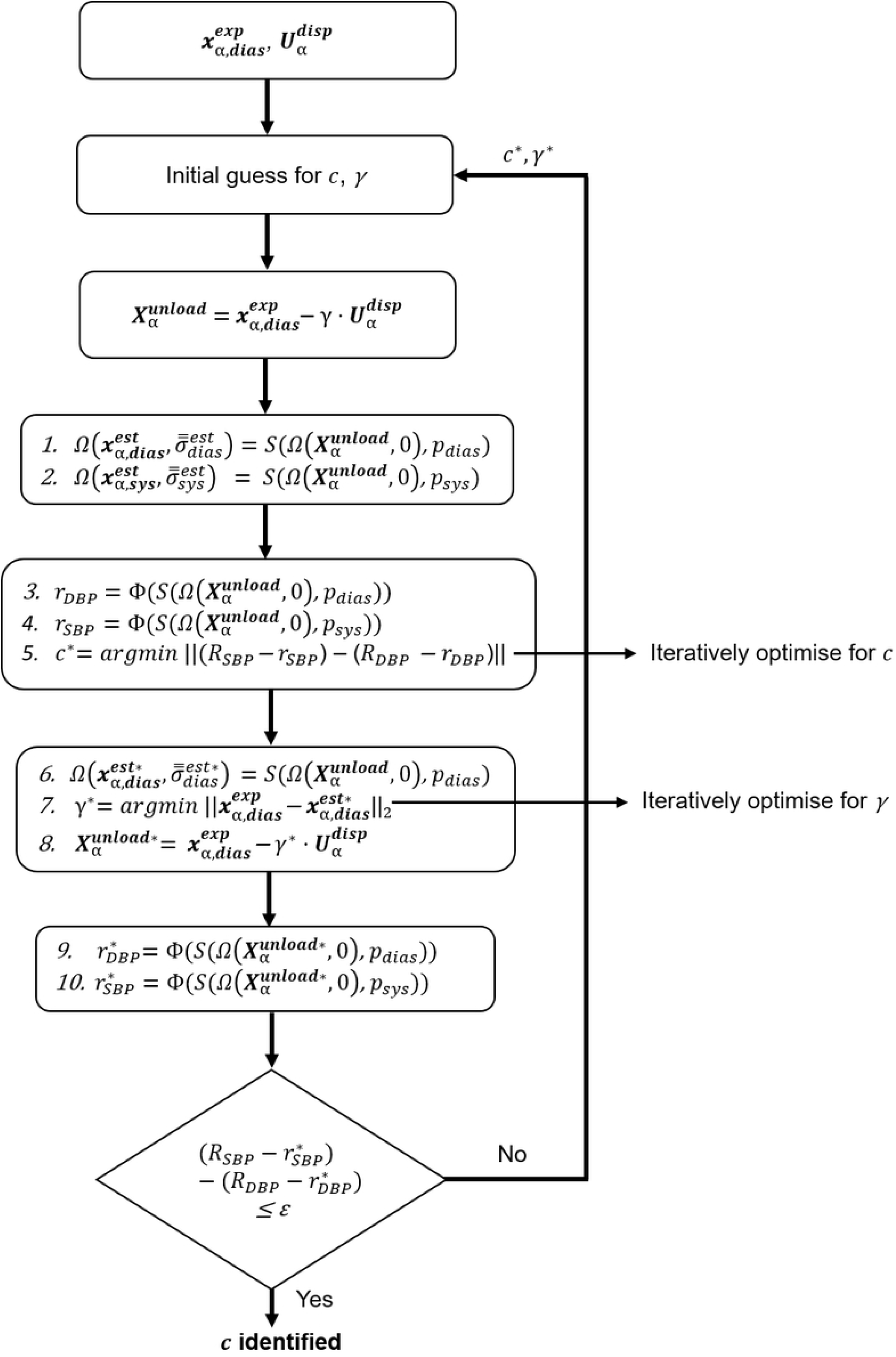
Workflow of the algorithm depicting the iterative process of estimating the mechanical parameter *c* and scaling factor *γ*. The subscript *α* in 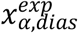 and 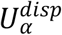, depict the type of geometry under consideration. In this paper *α* = cylinder and aorta.

### Finite element analysis

The finite element model is governed by the balance of linear momentum applied to the computational domain *Ω* (thus either the cylinder or aorta) and a constitutive model of the vessel wall.

The balance of linear momentum in the absence of body forces, and after neglecting inertial effects, reads:

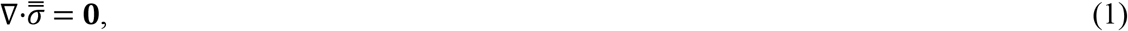

where, ∇ is the gradient operator and 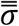 is the second-order Cauchy stress tensor.

To render the solution of the balance of linear momentum possible, boundary conditions (Dirichlet and Neumann) need to be satisfied where applicable.

Dirichlet boundary conditions are given by:

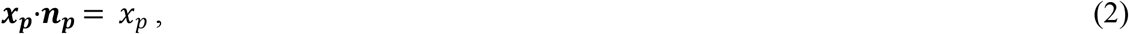

where, ***x***_***p***_ represents the prescribed nodal displacements in the direction ***n***_***p***_, on surfaces *Γ*_*d*_ (Fig 1-2). Neumann boundary conditions are given by:

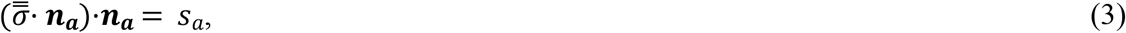

where, *s*_*a*_ denotes magnitude of applied traction in the direction ***n***_***a***_ on surfaces *Γ*_*n*_. As seen in Figs 1-2, the Neumann boundary condition in this study comprises only of blood pressure (*p*) application on the luminal side of the vessel.

An uncoupled Ogden hyperelastic strain energy density (*Ψ*) formulation [11] was used to describe the mechanical behaviour of the vessel geometries. The strain energy density (*Ψ*) was represented as:

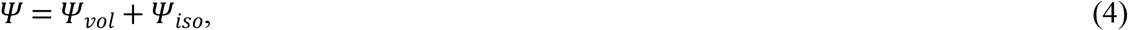

where,

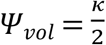 in(*J*)^2^ is the volumetric, and 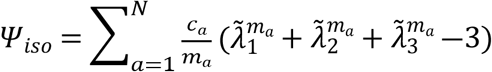 is the isochoric part of the strain energy density function. The parameters, *c*_*a*_ are shear modulus like mechanical parameters, parameter *κ* is the mechanical bulk modulus, while mechanical parameters *m*_*a*_ are unit-less and control the degree of nonlinearity. The parameter *N* defines the order of the Ogden model. Quantity *J* is the volume ratio defined by *J* = *λ*_1_*λ*_2_*λ*_3_. In Equation 4, 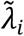 are the deviatoric principle stretches, defined by 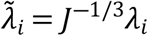. Here a second order form is used i.e. *N* = 2, with the following constraints: *c*_1_ = *c*_2_ = *c, m*_1_ = −*m*_2_ = *m* = 2, causing the model to reduce to a Mooney-Rivlin form with tension-compression symmetry [11]. To enforce nearly incompressible tissue behavior, the bulk modulus (*κ*) was constrained to *κ* = 100*c*.

Constitutively, the Cauchy stresses are derived by:

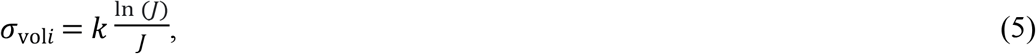

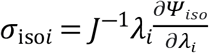 (*λ*_*i*_ are stretches in principal directions, and *i* = 1,2,3 represent three principal directions).

The mechanical equilibrium condition is said to be reached when balance of linear momentum is satisfied (Eq 1). The equilibrium solution resulting in the deformed shape and the stresses in the elements were obtained using FEBio’s [12] structural solver. The structural solver is denoted by *S* (Fig 1).

### Mechanical parameter estimation

This section provides a general description of the implementation of the data used for fitting and the mechanical parameter estimation algorithm particularised for aortic geometries. The aortic geometries are obtained from *in vivo* medical images.

### Experimental data used for fitting

In most cases, imaging data for the aortic geometries is obtained only for two pressure conditions, namely diastolic and systolic pressures. Henceforth, in this study, only the radii of the aortic geometry at diastolic and systolic pressures (*R*_*DBP*_ and *R*_*SBP*_ respectively) are used to assess the accuracy of our fitting procedure.

### Algorithm description

The algorithm is commenced by first assigning initial guesses for mechanical parameter *c* (cf. Eq. 4) and scaling parameter *γ*. Second, the image-extracted geometry at diastolic pressure, 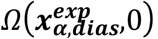, is pressurised (subscript ***α*** denotes the geometry under consideration) to systolic pressure:

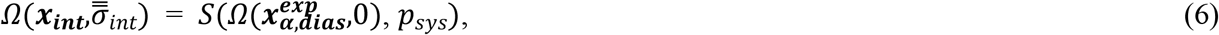

which results in the “intermediate” configuration 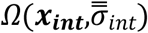. However, note that this intermediate configuration was obtained by assuming zero pre-stresses in the image-extracted geometry and thus not the correct unloaded configuration.

A displacement field is obtained by subtracting the nodes of the initial diastolic geometry from the intermediate configuration 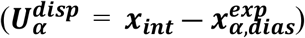. To consider the pre-stresses that are present in the diastolic geometry, an estimation of the unloaded configuration is made in the next step of the algorithm by subtracting a scaled displacement field from the initial diastolic configuration:

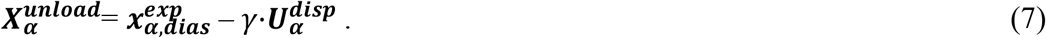

In which *γ* is an unknown scaling factor that needs to be estimated in addition to the mechanical parameter.

Hence, the algorithm was designed with a two-staged approach. The first-stage was to estimate the value of *c* by assuming that the assigned *γ* is the correct value. The second-stage comprised of utilising the newly estimated *c* to estimate *γ*.

For the initially assumed value of *c*, the unloaded configuration is then pressurised to diastolic and systolic pressures to obtain the deformed configuration for the respective pressures (steps 1 and 2; Fig. 3):

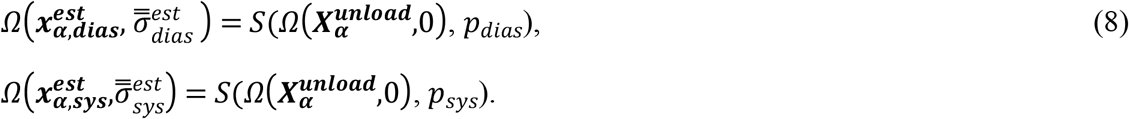

The inner radii of the diastolic and systolic geometries, *r*_*DBP*_ and *r*_*SBP*_ respectively, are calculated as the mean of distances from the luminal nodes to the centre (mean of nodal coordinates) of the lumen under consideration. The operator *Φ* (Fig. 3: Steps 3 and 4; Eq. 9) determines the procedure to extract the inner radii.

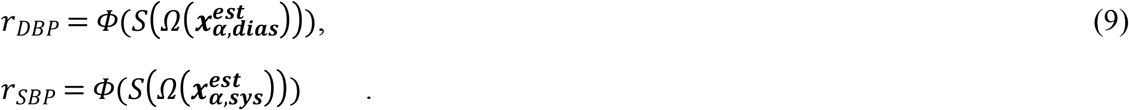

The first-stage optimisation is based on minimising an objective function (Eq.10) comprising of the differences between the experimental internal radii of diastolic and systolic geometries, *R*_*DBP*_ and *R*_*SBP*_, and *r*_*DBP*_ and *r*_*SBP*_ (Step 5; Fig. 3) by changing mechanical parameter *c*. Optimisation of the objective function leads to a new value of the mechanical parameter *c*^*^:

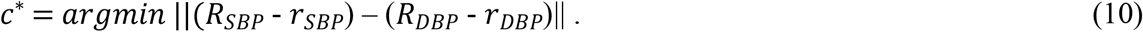

Next, the diastolic configuration 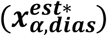 is produced by pressurising the unloaded configuration with *c*^*^ as the new mechanical parameter (Fig. 3 Step 6):

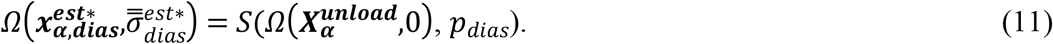

The second-stage optimisation is based on minimising an objective function (Eq.12) comprising of L^2^ – norm of the nodal differences of the experimental diastolic FE geometry 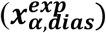 and the diastolic geometry 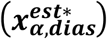 produced using the new mechanical parameter *c*^*^· The L^2^–norm is defined as 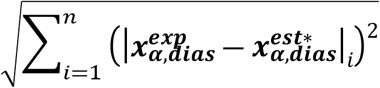 for all nodes = 1,2…*n*; and represented as 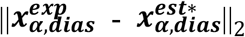. New scaling factor *γ*^*^ was determined as an outcome of the second-stage optimisation (Steps 6 and 7; Fig. 3):

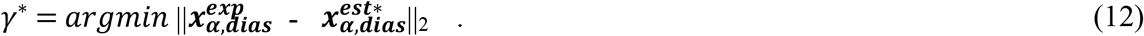

Subsequently, a new unloaded configuration is formed using *γ*^*^ (Step 8; Fig. 3)

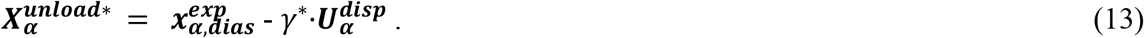

Furthermore, the new unloaded configuration is pressurised to obtain the internal radii at diastolic and systolic pressures (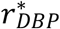 and 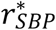; Fig. 3: steps 9-10):

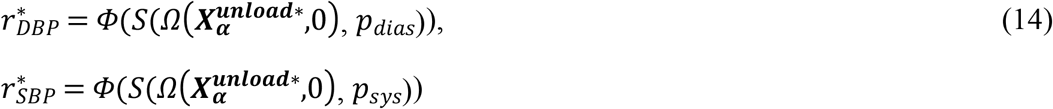

The differences between the experimental internal radii (*R*_*DBP*_ and *R*_*SBP*_) and the newly obtained internal radii (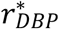 and 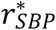) is compared by setting a tolerance value (ε) 0.01 mm. The decision to continue *DBP SBP* the iterations to estimate *c* and *γ* continues until the tolerance value is met (Fig. 3). The parameters c and *γ* are optimised in a least squares sense using the MATLAB function lsqnonlin which is configured to use the Levenberg-Marquardt algorithm [13].

### Method validation

The goal of validating the algorithm entailed determining the unloaded configuration and estimating the true mechanical parameter (c). Since the unloaded configuration and the true mechanical parameter cannot be obtained *in vivo*, reference data was generated synthetically to validate the algorithm.

#### Synthetically created reference data

Though reference data was generated for each geometry type, the method of generating the reference data for each geometry type differed slightly. The next two sections will delineate the methods to generate reference data (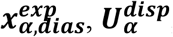, *R*_*DBP*_, and *R SBP*; cf. Fig. 3) for validating the algorithm.

#### Cylindrical geometry reference data

After creation of the unloaded geometry 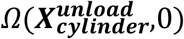 (Sec. *Cylindrical geometry*; Fig. 4), it was initially pressurised to a diastolic pressure (*p*_*dias*_ = 80 mmHg) to obtain the diastolic geometry 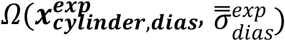 (Eq. 15; Fig. 4). Subsequently, the unloaded geometry was also pressurised to a systolic pressure (*p*_*sys*_ = 130 mmHg), hereby obtaining the systolic geometry 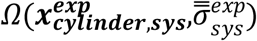 (Eq. 16; Fig. 4). The transformations to diastolic and systolic geometries are shown by the following equations:

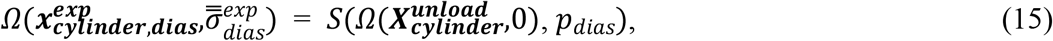

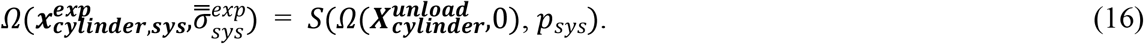

**Fig 4.**
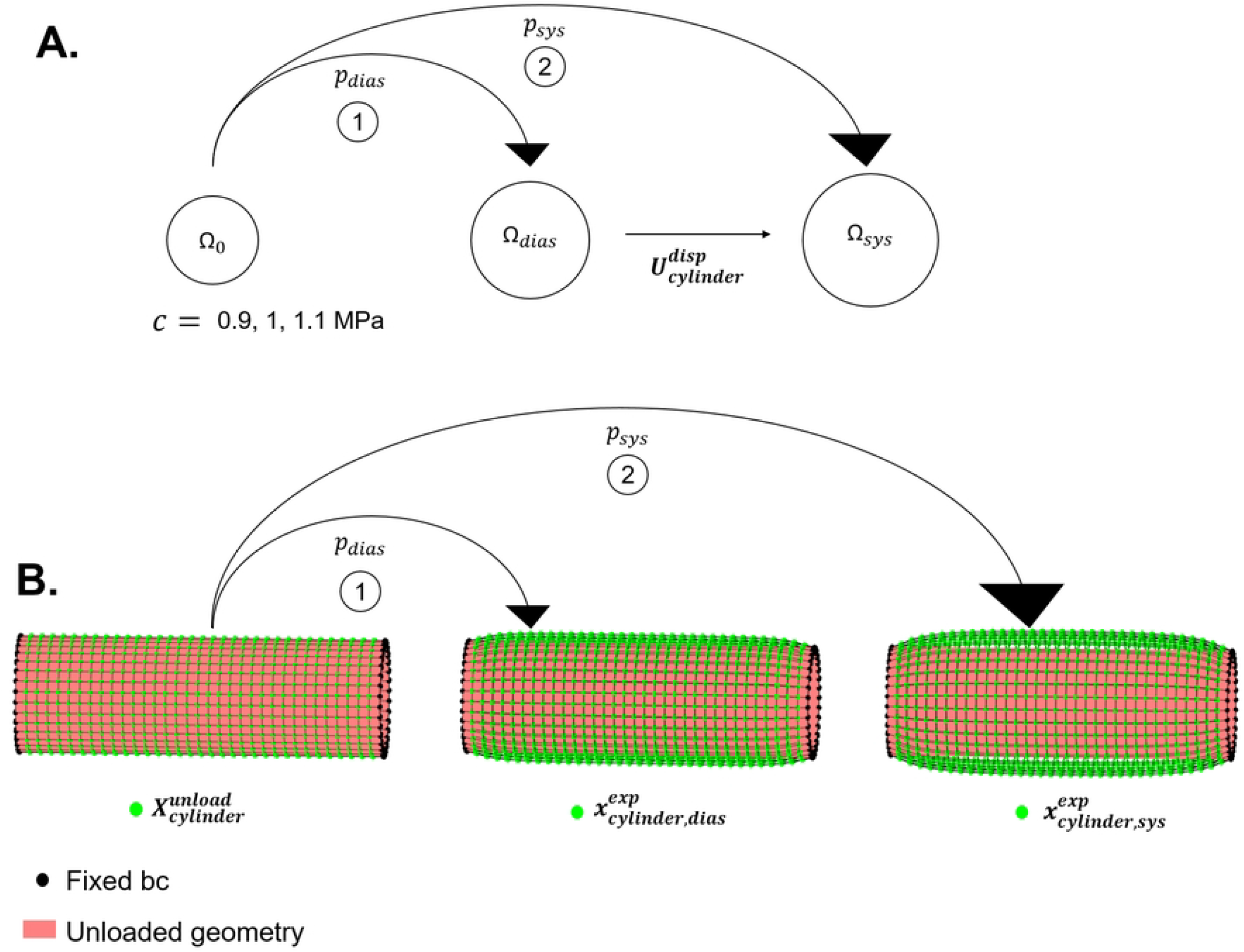
Steps involved to generate reference data for cylindrical geometry for a given unloaded configuration. (A): Schematic representation of the unloaded configuration (Ω_0_), diastolic configuration (Ω_*dias*_), and systolic configuration (Ω_*sys*_). (B): Finite element simulations depicting the nodes in unloaded 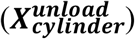 diastolic 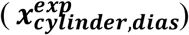 and systolic configurations 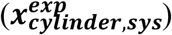 associated with Ω_0_, Ω_*dias*_, and Ω_*sys*_.

The displacement field 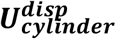 was calculated to be used as an input for mechanical parameter estimation

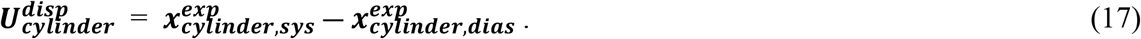

Additionally, *R*_*DBP*_ and *R*_*SBP*_ were created from nodes at the central inner boundary of the cylinder using:

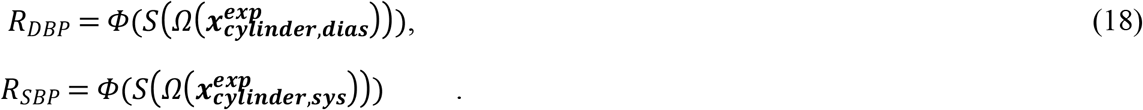

The reference data was generated for three different values of the shear modulus like mechanical parameter (*c*; cf. Eq. 4) – 0.9 MPa, 1 MPa and 1.1 MPa (Fig. 4).

#### Aortic geometry reference data

Generation of the reference data for the aortic geometry differed slightly from the cylindrical geometry. The initial aortic geometry *Ω*(***X***^***MRI***^,0) (Sec. *Image-based aorta geometry*; Ω_*MRI*_ in Fig. 5) was created from an MRI which was already at diastolic pressure, as opposed to the cylindrical geometry which was created with the unloaded configuration as the initial state. Therefore, an additional operation had to be performed on the initial aortic geometry to create an unloaded configuration (Steps 1-2 Fig. 5). The initial aortic geometry was pressurised (with zero displacement at root level) with a systolic pressure (*p*_*sys*_ = 130 mmHg) to obtain an intermediate configuration 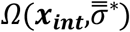 (Eq. 19; Ω_*int*_ in Fig. 5). The resulting displacement field 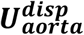 was computed (Eq. 20) and then scaled by a dimensionless scaling factor *γ* = 0·3. The value of *γ* was chosen arbitrarily. However, it was chosen in a way that the geometry would be prevented from scaling down such that the internal surface nodes would intersect, which would result in numerical errors. The unloaded aortic configuration 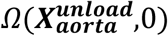 was then created (Eq. 21; Fig. 5) by scaling the obtained displacement field from the initial aortic geometry *Ω*(***X***^***MRI***^,0).

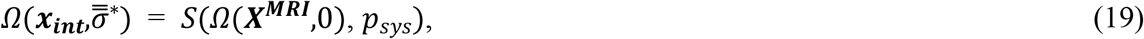

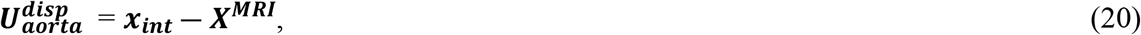

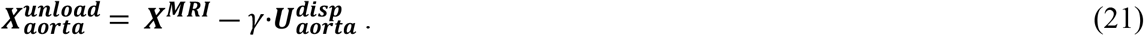

**Fig 5.**
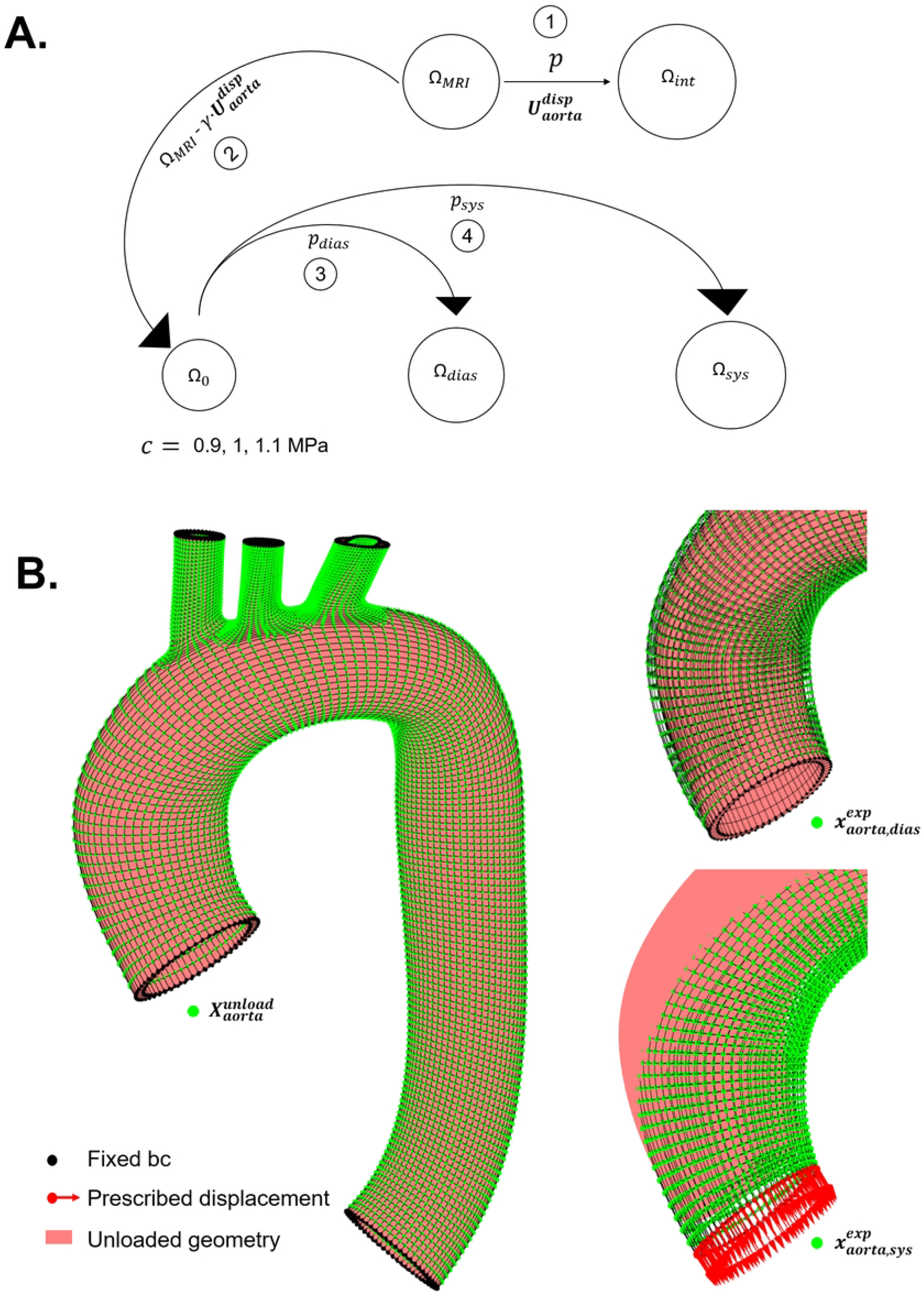
(A): Four-step procedure to generate reference data for aortic geometry: (i) initial MRI geometry Ω_*MRI*_ is inflated (till systolic pressure) to obtain an intermediate geometry Ω_*int*_, (ii) unloaded configuration Ω_0_ is obtained by subtracting the scaled nodal displacements) from MRI geometry Ω_*MRI*_, (iii) Subsequently pressurising Ω_0_ with diastolic and systolic pressures resulted in Ω_*dias*_, and Ω_*sys*_. (B Finite element simulations depicting the nodes in unloaded 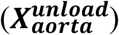 diastolic 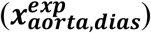 and systolic configurations 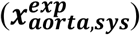 associated with Ω_0_, Ω*dias*, and Ω*sys*.

Diastolic and systolic configurations - 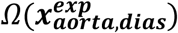 (Fig. 2B; Fig. 5) and 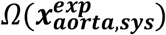 (Fig. Fig. 5), were generated in a similar way as the cylindrical geometry (Fig. 5; Eq. 22-23) for three values of shear modulus like mechanical parameter (*c* = 0.9 MPa, 1 MPa and 1.1 MPa; with same values of *p*_*dias*_ and *p*_*sys*_),

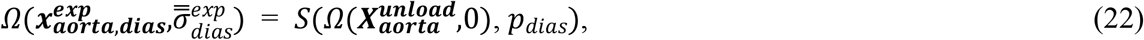

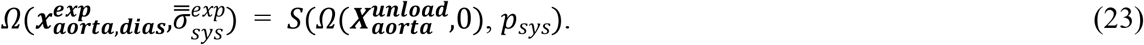

Contrary to the cylindrical geometry, displacement obtained from LVOT slice cine images was applied at *Γ*_*root*_ boundary of the aorta (Fig. 2) while pressurising the unloaded geometry to systolic pressure. The magnitude of the displacement was 8.7 mm. The displacement was not in the direction of the normal to *Γ*_*root*_. For the aortic geometry, *R*_*DBP*_ and *R*_*SBP*_ (Eq. 18) were created from the nodes located on the ascending aortic inner boundary (Fig. 2A).

### Simulations and analyses

#### Assessment of algorithm accuracy

On completion of the optimisation, the deviation (*Δ*c) of the estimated parameter *c* from the true value (reference value of *c*) was calculated as the absolute mean difference between the estimated value and the reference value. Accuracy (ζ) of the estimation, for each reference case, was measured by:

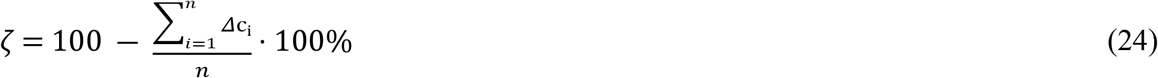

where, *n* is the number of initial guesses (in this case *n* = 4), and *Δ*c_i_ is the deviation for each estimation for a given value of reference value of *c*. The function tolerance for lsqnonlin was set to the default value provided by MATLAB which is 10^−6^.

#### Pre-stresses

The pre-stresses arising in the diastolic or systolic configuration are due to the deformations experienced by the geometry with respect to the unloaded configuration. Therefore, if the unloaded configuration is accurately estimated (along with mechanical parameter *c*), then the pre-stresses arising in the pressurised configurations would also be accurate. Therefore, for the aortic geometry, the effect of pre-stresses was indirectly determined by comparing estimated value of *γ* with the value of *γ* used to create the reference data.

However, since the cylindrical unloaded geometry was directly created as opposed to the aortic unloaded geometry, *γ* was not pre-assigned. Therefore, the pre-stresses in the diastolic configuration after estimation compared to the pre-stresses in the reference diastolic cylindrical geometries (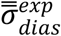 in Eq. 15). For comparison, the differences of L^2^–norms of the principal stresses of the reference diastolic configuration geometry 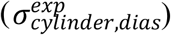 and the estimated diastolic configuration 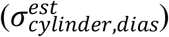 were calculated using:

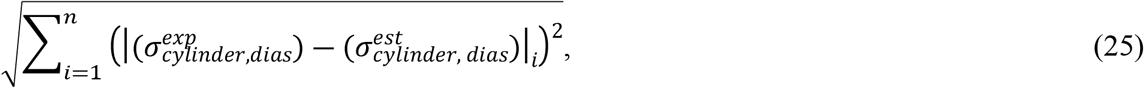

*i* represents the element number

#### Effect of different initial conditions

The algorithm was tested on four different sets of initials guesses for *c* and *γ* required to initiate the parameter estimation process (Fig. 5) for the cylindrical and aortic reference data. The four sets of initial guesses for *c* and *γ* were – (i) 0.5 MPa and 1, (ii) 0.75 MPa and 0.75, (iii) 1 MPa and 0.5, and (iv) 1.5 MPa and 0.25.

The initial values for *c* were chosen to check if convergence to the true value was obtained when the material initially behaved flexibly (*c* = 0.5 MPa) or stiffly (*c* = 1.5 MPa). The values of 0.75 MPa and 1 MPa were chosen to check the convergence with the middle value of the stiffness range (0.5 – 1.5 MPa). The initial values of *γ* are lying within the range as previously recommended [8].

To check the convergence for *γ* to the true value for aortic reference data (0.3; cf. Sec. *Aortic geometry reference data*), 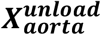 in the algorithm (Fig. 3) was defined as 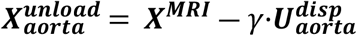 instead of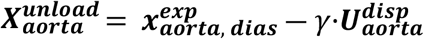., since the unloaded configuration for the aorta was created from ***X***^***MRI***^ (Eq. 21) and not from 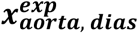.

## Results

Table 1 describes the outcome of the estimated parameter values for the cylindrical and aortic geometries. It can be observed that the mechanical parameter *c* was slightly overestimated for the cylinder, while underestimated for the aorta. However, independent from the initial guesses, the estimations resulted in acceptable accuracies of *c* for the cylindrical (98.43%, 98.50%, 98.50%), and the aortic (99.15%, 99.22%, 99.20%) geometries. The differences in stresses for cylindrical geometry between the reference data and post-estimation of mechanical parameters ranged from 0.005 MPa to 0.007 MPa. The average values of *γ* for the cylindrical geometry were 1.279, 1.313 and 1.340, whereas the aortic geometry resulted in average values of 0.304, 0.304, and 0.303 for reference *c* values of 0.9, 1 and 1.1 MPa respectively.

**Table 1.** Initial guesses and estimated outcomes of *c* and *γ* parameters for reference values of *c* for cylindrical and aortic geometries.

| Geometry | Reference $c$ (MPa) | Initial guesses | | Estimated values | | $\Delta c$ (MPa) | Accuracy ($\zeta$) |
| --- | --- | --- | --- | --- | --- | --- | --- |
| | | $c$ (MPa) | $\gamma$ | $c$ (MPa) | $\gamma$ | | |
| Cylinder | 0.9 | 0.5 | 1 | 0.930 | 1.260 | 0.030 | 98.43% |
|  |  | 0.75 | 0.75 | 0.908 | 1.289 | 0.008 |  |
|  |  | 1 | 0.5 | 0.911 | 1.285 | 0.011 |  |
|  |  | 1.5 | 0.25 | 0.914 | 1.281 | 0.014 |  |
|  | 1 | 0.5 | 1 | 1.032 | 1.293 | 0.032 | 98.50% |
|  |  | 0.75 | 0.75 | 1.008 | 1.322 | 0.008 |  |
|  |  | 1 | 0.5 | 1.010 | 1.320 | 0.010 |  |
|  |  | 1.5 | 0.25 | 1.010 | 1.317 | 0.010 |  |
|  | 1.1 | 0.5 | 1 | 1.134 | 1.320 | 0.034 | 98.50% |
|  |  | 0.75 | 0.75 | 1.107 | 1.350 | 0.007 |  |
|  |  | 1 | 0.5 | 1.109 | 1.347 | 0.009 |  |
|  |  | 1.5 | 0.25 | 1.110 | 1.345 | 0.010 |  |
| Aorta | 0.9 | 0.5 | 1 | 0.895 | 0.303 | 0.005 | 99.15% |
|  |  | 0.75 | 0.75 | 0.897 | 0.302 | 0.003 |  |
|  |  | 1 | 0.5 | 0.879 | 0.313 | 0.021 |  |
|  |  | 1.5 | 0.25 | 0.905 | 0.297 | 0.005 |  |
|  | 1 | 0.5 | 1 | 0.996 | 0.302 | 0.004 | 99.22% |
|  |  | 0.75 | 0.75 | 0.997 | 0.302 | 0.003 |  |
|  |  | 1 | 0.5 | 0.979 | 0.312 | 0.021 |  |
|  |  | 1.5 | 0.25 | 1.003 | 0.298 | 0.003 |  |
|  | 1.1 | 0.5 | 1 | 1.096 | 0.302 | 0.004 | 99.20% |
|  |  | 0.75 | 0.75 | 1.098 | 0.301 | 0.002 |  |
|  |  | 1 | 0.5 | 1.079 | 0.311 | 0.021 |  |
|  |  | 1.5 | 0.25 | 1.105 | 0.297 | 0.005 |  |

## Discussion

In the current study, we proposed a method to estimate mechanical parameters of the ascending aorta using an inverse finite element analysis formulation while considering the *in vivo* pre-stresses. The presented method utilises a simplistic two-staged approach to estimate the mechanical parameters. The two-staged approach enabled to determine the unloaded configuration as well, which is physiologically not feasible, yet important for considering the pre-stresses in the artery while estimating the mechanical parameters. Our method is based on previously proposed methods to determine the unloaded configurations of blood vessels [5, 8], with an additional ability to estimate mechanical parameters by considering the pre-stresses [6, 9]. Though the backward incremental method has proven effective for mechanical characterisation of arteries [6, 9], it does not calculate the pre-stresses or the unloaded configuration directly but rather calculates an equilibrium configuration by incrementally applying *in vivo* pressure to the *in vivo* geometry. During incrementally pressurising the *in vivo* geometry, the deformations were discarded. The resulting effect is an *in vivo* geometry with pre-stresses only. However, to perform the backward incremental method, the structural solver to perform finite element analysis needs to be adjusted in such a way that on incrementally pressurising a given arterial geometry, only the stress field is updated after each iteration while the geometry is not updated based on the deformations [5]. Our method warrants the use of finite element solvers without creating any alterations in the source code of the structural solver to consider the effect of pre-stresses present under *in vivo* conditions. The difference between our method and the backward incremental lies in the fact that our method utilises the unloaded configuration explicitly to calculate the stresses in the *in vivo* configuration, while the backward incremental method has an implicit consideration of the unloaded configuration while explicitly solving for *in vivo* pre-stresses [5, 6].

To validate the utility of our method, the algorithm was first implemented on a cylindrical case. The cylindrical case was chosen in order to simplify the deformation field to strictly radial direction expansion. To achieve radial direction expansion only, our cylinder case with fixed ends was chosen to be long in order to ameliorate the effect of boundary conditions. Nevertheless, the geometrical features (radius and thickness) of the cylinder and the pressure conditions it was subjected to were similar to a normal ascending aorta. Accuracy checks were performed on the estimated mechanical parameter *c* by comparing it with the true value with which the reference data was produced synthetically. The minimum accuracy of the estimated *c* for the cylindrical geometry was 98.43%.

Following the validation of the cylindrical case, the method was implemented on the aortic geometry. The boundary conditions for the aortic geometry were chosen in order to mimic the *in vivo* conditions. However, since the displacement of the aortic root cannot be obtained from the *in vivo* unloaded configuration to the diastolic configuration, zero-displacement boundary condition of the aortic root was chosen while pressurising the unloaded configuration to the diastolic pressure. Therefore, the aorta did not experience any axial stretch while deforming from unloaded configuration to the configuration at the diastolic pressure. Unavailability of the *in vivo* root displacement information from the unloaded configuration to the diastolic configuration poses as a limitation of this method, which may affect the accuracy of the mechanical parameter *c*.

Moreover, since the value of aortic root displacement at systolic pressure with respect to the diastolic geometry could be obtained from MRI, aortic root displacement was prescribed at the root level while pressurising the geometry from unloaded configuration to the systolic pressure. Applying root displacement to the directly unloaded configuration while pressurising to the systolic pressure will not cause a difference in the stresses in the vessel wall when compared to pressurising the unloaded geometry to the diastolic pressure first with fixed root, and then pressurising the diastolic configuration till the systolic pressure with root displacement. This is because the deformations at the final state (systolic pressure) are independent of the deformation path for hyperplastic materials.

Application of zero-displacement boundary condition at the root level while pressurising to diastolic pressure and a prescribed boundary while pressurising to the systolic pressure is not physiological, since the displacement of the root would be occurring at both diastolic and systolic geometry. Additionally, the displacement field 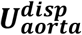, used to calculate the unloaded aortic geometry (Eq. 20) was determined by applying a positive (systolic) pressure to the geometry created from MRI. It is to be noted that 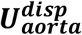 might not be a physiological deformation field that could be existing if the blood pressure was removed from the aorta. Nonetheless, the minimum accuracy of the estimated *c* for the aortic geometry was 99.15% regardless of the initial guesses for *c* in our algorithm.

Regardless of the fact that precision on boundary conditions and displacement fields would provide more accurate results in terms of stresses and strains in the geometry under consideration, the goal of this study was to develop a simplistic method to estimate aortic mechanical parameters. Similar line of reasoning was also implemented while considering a Mooney-Rivlin type of formulation as the mechanical property of the cylindrical and the aortic geometry. Even though the aorta does not exhibit isotropic behaviour [4], an isotropic formulation was implemented in this study due to the scarcity of available data (which is usually the case in clinical settings) to fit for a complex mechanical law exhibiting an anisotropic and exponential behaviour [4]. Nevertheless, the method proposed in this study could be used for complex mechanical behaviours exhibiting anisotropy, given that sufficient experimental data (in terms of multiple geometries at multiple pressure conditions) from the medical images is available.

Despite the scarcity of data, our method can be implemented (using isotropic formulation) to identify the differences in mechanical properties between healthy and diseased groups.

## Conclusion

In conclusion, the method presented in this paper demonstrates the ability to estimate mechanical parameters of an ascending aorta with admissible accuracy (error < 2%), by preserving pre-stresses in the aortic wall.

## Data availability

The authors ensure the availability of the MATLAB codes utilised in this study to any interested researcher. The request for the codes must be sent to the corresponding author.

## References

1. Choudhury N, Bouchot O, Rouleau L, Tremblay D, Cartier R, Butany J, et al. Local mechanical and structural properties of healthy and diseased human ascending aorta tissue. Cardiovasc Pathol. 2009;18(2):83–91. Epub 2008/04/12. doi: 10.1016/j.carpath.2008.01.001. PubMed PMID: 18402840.

2. Vorp DA, Schiro BJ, Ehrlich MP, Juvonen TS, Ergin MA, Griffith BP. Effect of aneurysm on the tensile strength and biomechanical behavior of the ascending thoracic aorta. Ann Thorac Surg. 2003;75(4):1210–4. Epub 2003/04/10. doi: 10.1016/s0003-4975(02)04711-2. PubMed PMID: 12683565.

3. Johansson G, Markström U, Swedenborg J. Ruptured thoracic aortic aneurysms: a study of incidence and mortality rates. Journal of vascular surgery. 1995;21(6):985–8.

4. Humphrey JD. Cardiovascular solid mechanics: cells, tissues, and organs: Springer Science & Business Media; 2013.

5. Bols J, Degroote J, Trachet B, Verhegghe B, Segers P, Vierendeels J. A computational method to assess the in vivo stresses and unloaded configuration of patient-specific blood vessels. Journal of computational and Applied mathematics. 2013;246:10–7.

6. De Putter S, Wolters B, Rutten M, Breeuwer M, Gerritsen F, Van de Vosse F. Patient-specific initial wall stress in abdominal aortic aneurysms with a backward incremental method. Journal of biomechanics. 2007;40(5):1081–90.

7. Chen H, Wang W, Chen Q, Li Z. Numerical Determination of the Circumferential Residual Stress of Porcine Aorta by Pulling-Back Method. Acta Mechanica Solida Sinica. 2021;34(3):346–55.

8. Raghavan M, Ma B, Fillinger M. Non-invasive determination of zero-pressure geometry of arterial aneurysms. Annals of biomedical engineering. 2006;34(9):1414–9.

9. Van Disseldorp E, Petterson N, Rutten M, Van De Vosse F, van Sambeek M, Lopata R. Patient specific wall stress analysis and mechanical characterization of abdominal aortic aneurysms using 4D ultrasound. European Journal of Vascular and Endovascular Surgery. 2016;52(5):635–42.

10. Moerman KM. GIBBON: the geometry and image-based bioengineering add-on. Journal of Open Source Software. 2018;3(22):506.

11. Moerman KM, Simms CK, Nagel T. Control of tension–compression asymmetry in Ogden hyperelasticity with application to soft tissue modelling. Journal of the mechanical behavior of biomedical materials. 2016;56:218–28.

12. Maas SA, Ellis BJ, Ateshian GA, Weiss JA. FEBio: finite elements for biomechanics. Journal of biomechanical engineering. 2012;134(1).

13. Marquardt DW. An algorithm for least-squares estimation of nonlinear parameters. Journal of the society for Industrial and Applied Mathematics. 1963;11(2):431–41.

